# Lysine-36 of *Drosophila* histone H3.3 supports adult longevity

**DOI:** 10.1101/2023.09.28.559962

**Authors:** John C. Brown, Benjamin D. McMichael, Vasudha Vandadi, Aadit Mukherjee, Harmony R. Salzler, A. Gregory Matera

## Abstract

Aging is a multifactorial process that disturbs homeostasis, increases disease susceptibility, and ultimately results in death. Although the definitive set of molecular mechanisms responsible for aging remain to be discovered, epigenetic change over time is proving to be a promising piece of the puzzle. Several posttranslational histone modifications (PTMs) have been linked to the maintenance of longevity. Here, we focus on lysine-36 of the replication-independent histone protein, H3.3 (H3.3K36). To interrogate the role of this residue in *Drosophila* developmental gene regulation, we generated a lysine to arginine mutant that blocks the activity of its cognate modifying enzymes. We found that an *H3.3B^K36R^* mutation causes a significant reduction in adult lifespan, accompanied by dysregulation of the genomic and transcriptomic architecture. Transgenic co-expression of wild-type *H3.3B* completely rescues the longevity defect. Because H3.3 is known to accumulate in non-dividing tissues, we carried out transcriptome profiling of young vs aged adult fly heads. The data show that loss of H3.3K36 results in age-dependent misexpression of NF-κB and other innate immune target genes, as well as defects in silencing of heterochromatin. We propose H3.3K36 maintains the postmitotic epigenomic landscape, supporting longevity by regulating both pericentric and telomeric retrotransposons and by suppressing aberrant immune signaling.

## Introduction

Eukaryotic genomes are condensed in the nucleus of each cell as chromatin, a dynamic blend of histones, proteins and DNA. Nucleosomes, the fundamental unit of chromatin, comprise ∼150 base pairs of DNA wrapped around an octamer of histones. Nucleosomes not only allow organisms to compress and store vast amounts of genetic information, but they also serve as steric barriers to a variety of DNA-templated processes [1–3]. Accordingly, molecular mechanisms have evolved to alter and regulate genome accessibility. Included among these mechanisms for chromatin remodeling are the covalent post-translational modification (PTM) of histones and the incorporation of histone variants [4–6].

Histone tail residues sustain a remarkable diversity of PTMs, including methylation, acetylation, ubiquitylation, sumoylation, and phosphorylation [7, 8]. Histone PTMs are hypothesized to alter local chromatin states via direct and indirect methods. For example, lysine acetylation alters chromatin directly by neutralizing positively charged lysine residues and weakening electrostatic interactions between histones and DNA [9]. In contrast, other modifications function through recruitment of effector proteins [8].

The incorporation of variant histones further increases regulatory complexity. In *Drosophila*, the histone gene complex (*HisC*) encodes all five canonical histone proteins. It is comprised by a cluster of ∼100 tandemly repeated, 5kb repeat units that encode H2A, H2B, H3, H4, as well as the linker histone H1. Canonical (replication-dependent) histones are synthesized and incorporated into chromatin during S phase of the cell cycle. Variant (replication-independent) histones are encoded by genes located outside of *HisC* and are synthesized and incorporated independently of DNA replication. Variant histones typically contain a few amino acid residue changes from their canonical counterparts, which can alter their PTMs, stability within the nucleosome, and interactions with other chromatin components [6].

Here, we focus on lysine-36 of the replication-independent histone, H3.3, and its role in regulating lifespan. H3.3 differs from the canonical H3 (H3.2) by four amino acids [10]. These few differences enable a number of unique protein-protein interactions and PTM preferences [11, 12]. Ascomycete genomes, including budding yeast and other fungi, possess only one non-centromeric H3 protein, which is orthologous to metazoan H3.3. In budding yeast, mutation of either H3K36 or Set2, the enzyme that catalyzes mono-, di-and tri-methylation of H3K36, causes replicative aging defects [13]. In nematodes, inactivation of met-1, the Set2 orthologue, also decreases lifespan, whereas loss of H3.3 decreases the lifespan of long-lived *C. elegans* mutants [14–16]. Although these results suggest that H3.3K36 plays a conserved role in metazoan lifespan regulation, direct genetic tests of a role for this histone residue in aging have been lacking.

To investigate the function of H3.3K36 in *Drosophila* gene expression regulation, we previously generated a lysine to arginine mutant, *H3.3B^K36R^* [17]. Here, we show that an *H3.3B^K36R^* mutation (expressed in an *H3.3A^null^* background) causes age-dependent dysregulation of the transcriptome, resulting in a significantly reduced adult lifespan. This longevity defect can be rescued by ectopic expression of a wild-type *H3.3B* transgene. We also show that loss of H3.3K36 histones causes overexpression of NF-κB target genes in the adult brain, a disruption in several key aging-related metabolic processes, and progressive deterioration in silencing of heterochromatin. Indeed, we found clear evidence for age-dependent upregulation of repetitive elements, particularly LTR-type retroposons.

Remarkably, expression of LINE-like elements specifically involved in telomere maintenance was decreased. These findings lead us to propose that H3.3K36 supports longevity not only by promoting cellular homeostasis, but also by suppressing the activity of transposable elements and innate immune response genes in the adult brain.

## Results

### *H3.3^K36R^* causes an adult lifespan defect

*Drosophila* histone H3.3 protein is encoded by two genes, *H3.3A* on the second chromosome and *H3.3B* on the X chromosome [18]. We previously engineered a missense mutation in *H3.3B* (*H3.3B^K36R^*) using Cas9-mediated homologous DNA repair, for details see [17]. By combining the *H3.3B^K36R^* mutation with a deletion of *H3.3A* (*H3.3A^null^*), we generated *H3.3^K36R^* animals that completely lack H3.3K36 (Fig. 1A). We also utilized an *H3.3B^WT^* transgene inserted onto the third chromosome [19] to generate *H3.3A^null-WTR^* and *H3.3^K36R-WTR^* (wild-type rescue) flies (Fig. 1A). As shown in Fig. 1B, the majority *H3.3A^null^, H3.3^K36R^*, and *H3.3^K36R-WTR^* animals complete development and eclose as adults at similar frequencies to a wild type Oregon R (Ore-R) control. Thus, arginine substitution of H3.3K36 has no significant effect on organismal viability.

**Figure 1.**
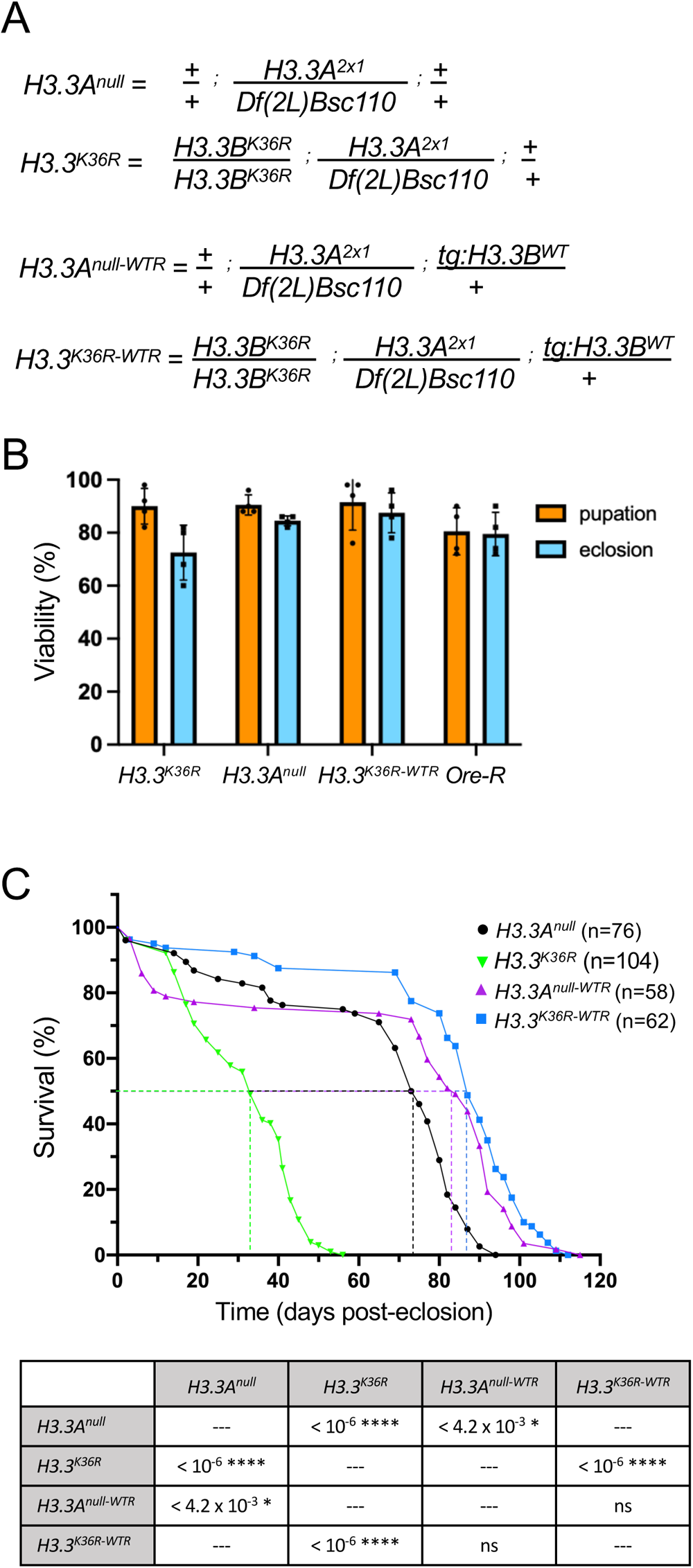
*H3.3^K36R^* mutation reduces adult lifespan relative to *H3.3A^null^* and wild-type rescue (WTR) controls. **(A)** Detailed genotypes for *H3.3A^null^*, *H3.3^K36R^*, *H3.3A^null-WTR^*, and *H3.3^K36R-WTR^* animals. The notation *tg:H3.3B^WT^* corresponds to the rescue transgene. **(B)** Developmental viability assay. For each genotype, pupation and eclosion frequencies (%) were calculated and plotted (50 larvae per biological replicate vial). The mean and SD of these percentages are shown. Oregon-R (Ore-R) was used as an additional control. **(C)** Adult longevity assays for *H3.3A^null^* and *H3.3A^null-WTR^* controls, and for *H3.3^K36R^* and *H3.3 ^K36R-WTR^* flies. Median lifespan was determined (dotted lines) by identifying the day at which 50% of the animals survived. Statistical comparison of survival curves using Gehan-Breslow-Wilcoxon tests are presented in the accompanying table. A Bonferroni correction for multiple comparisons was employed, resulting in the following adjusted significance values: * p<0.0125, ** p<0.0025, *** p<2.5 x 10^-4^, ****p<2.5 x 10^-5^.

To understand the role that this residue might play in longevity, we carried out adult lifespan assays. We measured the lifespan of *H3.3^K36R^* mutants and compared the results to those obtained for *H3.3A^null^* control animals (Fig. 1C, graph). Note that the *H3.3A^null^* mutation has a small, but significant decrease in median lifespan by comparison to an Ore-R wildtype strain (Fig. S1). As defined by the number of days (post-eclosion) at which 50% of the animals survived, the K36R mutation has a pronounced and highly significant effect on adult lifespan (Fig. 1C, table). Furthermore, ectopic expression of wildtype H3.3 protein fully rescued this defect. That is, the median survival time of *H3.3^K36R^* mutants was roughly 33 days post-eclosion, whereas that of *H3.3^K36R-WTR^* control animals was 87 days (Fig. 1C, graph). Thus, we conclude that mutation of H3.3 lysine-36 causes a substantial reduction in adult lifespan. Consistent with previous studies showing that H3.3 is essential for lifespan extension in nematodes [15], this work shows that lysine-36 plays a direct role in this process.

### Gene expression analysis in young vs. old fly heads from *H3.3^K36R^* mutants and controls

Organismal lifespan is influenced by multiple interconnected factors, many of which are unknown, and most of which are poorly understood [20–22]. Although cellular senescence is a natural and beneficial process in certain contexts, the accumulation of senescent cells over time is thought to contribute to aging and age-related diseases [23]. Despite their relatively short lifespan, many fundamental mechanisms underlying aging are conserved between fruit flies and humans, making *Drosophila* a valuable model for studying senescence-related processes. The brain of the adult fly is composed primarily of diploid postmitotic cells that rely on histone variant-specific assembly mechanisms for nucleosome turnover [24]. Indeed, H3.3 is reportedly enriched in the adult fly brain, and its deposition increases with age [25]. Thus, brain cells should be particularly sensitive to potential effects of a histone H3.3 missense mutation.

To gain insight into mechanisms by which H3.3K36 might influence age-dependent regulation of gene expression, we carried out transcriptome profiling of young vs. old fly heads. As detailed in the Materials and Methods, adult head RNA was extracted from six replicates (three male and three female) of each age:genotype pair (young and old *H3.3A^null^* and young and old *H3.3^K36R^* animals). Libraries were prepared from total, Ribo-minus RNA samples and subjected to 100nt, paired-end RNA-sequencing (RNA-seq) using standard procedures (see Methods for details). Principle component analysis of the dataset (Fig. 2A) shows that the samples segregate primarily on the basis of age (PC1) and sex (PC2). Furthermore, the old adult samples can be distinguished by genotype-specific differences (Fig. 2A).

**Figure 2.**
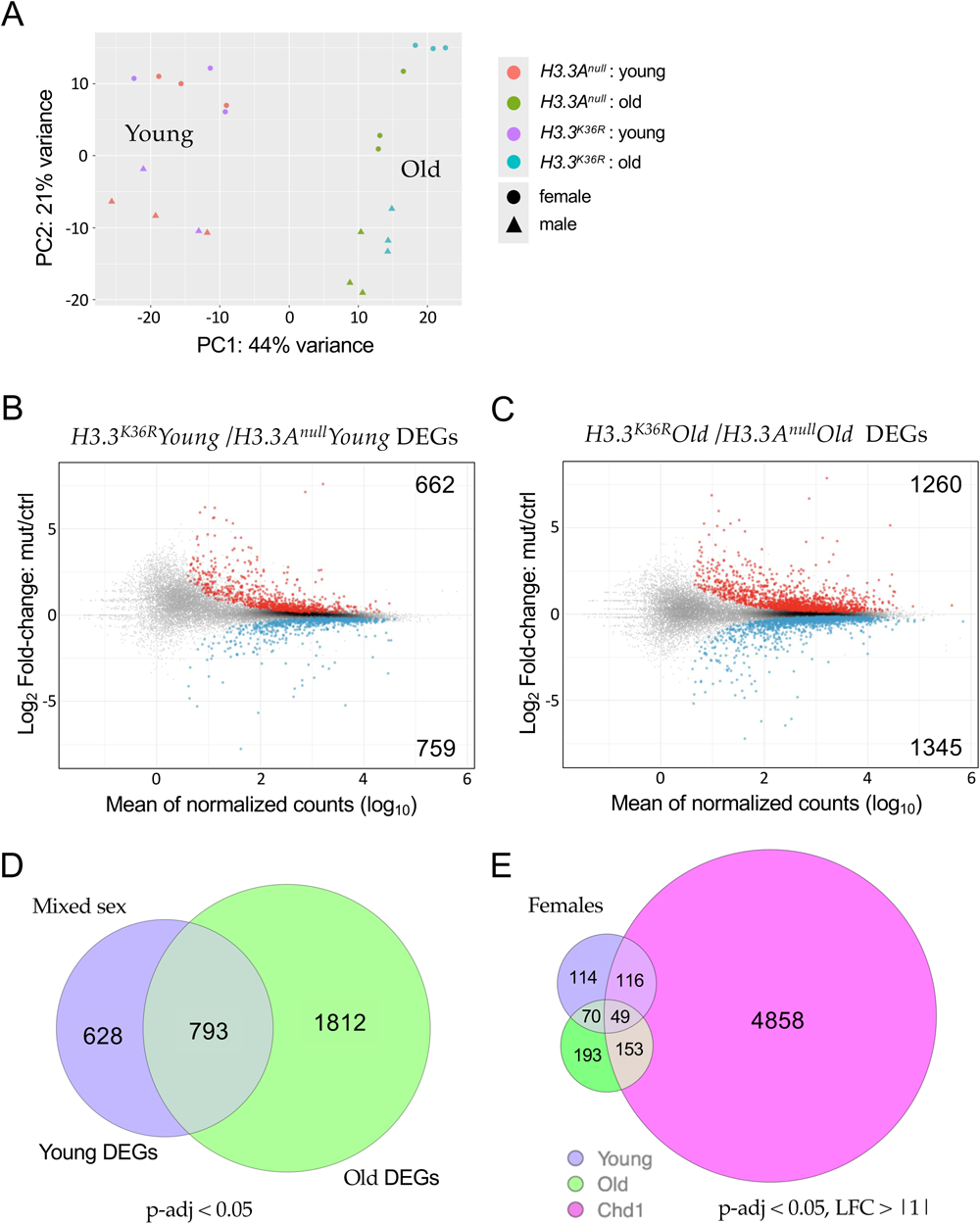
Transcriptomic profiling of young and old adult fly brains. **(A)** Principle Component Analysis (PCA) of young (0-2 d post-eclosion) and old (21-25 d) *H3.3A^null^* and *H3.3^K36R^* fly heads, labeled by sex. **(B)** M/A plot comparing gene expression from mixed sex *H3.3^K36R^* and *H3.3A^null^* young fly heads. Red and blue dots represent differentially expressed genes (DEGs) that were significantly (adjusted p-value, p-adj < 0.05) up-or down-regulated, respectively. The number of DEGs in each direction is shown in the upper and lower corners. **(C)** M/A plot of mixed sex *H3.3^K36R^* vs. *H3.3A^null^* transcripts from old adults. Labeling and significance as per panel B. **(D)** Venn diagram comparing overall number of DEGs (p-adj <0.05) between *H3.3A^null^* and *H3.3^K36R^* genotypes, young and old mixed sex fly heads. **(E)** Venn diagram of DEGs (p-adj < 0.05 and Log2 Fold-change ≥ |1|) from female *H3.3^K36R^* (young or old) versus aged (20 d) female *Chd1* fly heads. Data for *Chd1* provided by [25].

In order to identify differentially expressed genes (DEGs), we plotted the log2 fold-change (*H3.3^K36R^*/*H3.3A^null^*) for each transcript, versus the mean of normalized counts for that transcript. Among the younger flies, we found that there were 1,421 genes (662 up, 759 down) that were differentially expressed (p-adj < 0.05) in the mutant vs. control samples (Fig. 2B). A similar M/A plot comparing the older mutant and control animals showed that there were 2,605 DEGs (1260 up and 1345 down, Fig. 2C). Deposition of H3.3 protein in adult heads has been shown to increase with age from ∼17% of total H3 in newly-eclosed animals to ∼34% in 20-25 day old flies like the ones used here [25]. Consistent with this finding, we identified nearly twice as many DEGs in the older dataset, 793 of which overlapped with those in the younger animals (Fig. 2D).

To maximize the chances of capturing H3.3-specific effects, we focused our attention on DEGs identified in the older age group, and compared them to an RNA-seq dataset prepared from similarly-aged female *Chd1* (*chromodomain-helicase-DNA binding protein 1*) null mutants, using a separate DESeq2 model using data only from females [25]. Chd1 protein is part of a histone chaperone complex that functions together with Hira (Histone regulator A) to specifically deposit H3.3 into chromatin in the adult brain and other post-mitotic tissues [24]. *Chd1* null animals display a very similar adult longevity phenotype to *H3.3^K36R^* (K36R) mutants [25]. Differential expression analysis of *Chd1* null animals identified roughly 12,000 DEGs (p-adj < 0.05). Unsurprisingly, most of the previously identified (young + old) K36R DEGs were contained within this large set (see supplementary Figure S2A). We therefore applied more stringent cutoffs (p-adj < 0.05, log2 FC > |1|) to highlight the genes with the greatest magnitude changes in all three groups and found that the overlap was greatly reduced. For this more stringent comparison, we also restricted it to our female K36R datasets, because the Chd1 DEGs were derived from female-only fly heads (Fig. 2E). Strikingly, the degree of overlap between the old K36R and Chd1 DEGs went from roughly 84% to around 43% (Fig. S2B). Moreover, all of the non-Chd1 overlapping transcripts in the old female K36R dataset remained above the higher stringency threshold (Fig. 2E). Thus, the K36-specific DEGs, generally represented larger magnitude changes.

Interestingly, the female K36R fly head transcriptome appears to be affected earlier, and disrupted to a greater extent, than that of the male (Fig. 3A). Parsing out the longevity data in Fig. 1 by sex revealed that female K36R mutants had a median lifespan of around 30 d, compared to 40 d for males (Fig. S3). Functional enrichment analysis of the old female DEGs produced a number of senescence-associated gene ontology (GO) terms. These include genes involved in metabolic pathways that include cytochrome P450 and other enzymes, such as oxidoreductases (Fig. 3B). In contrast, GO analysis of old male K36R mutants identified detoxification factors, serine peptidases, and ecdysteroid insect hormone enzymes (Fig. 3B). Interestingly, brain extracellular region proteins were identified in both sexes. This category of genes is among the hallmarks of the senescence-associated secretory phenotype (SASP), which has been implicated as a driver of numerous age-related disorders [26–28]. Another prominent GO term category, identified in both males and females, centers around the oxidization and transport of glutathione (Fig. 3B). In studies from nematodes to mice, glutathione has been broadly implicated in brain disorders and cellular senescence [29–31]. Altogether, these data support the view that H3.3K36 is important for maintaining cellular homeostasis in the adult brain.

**Figure 3.**
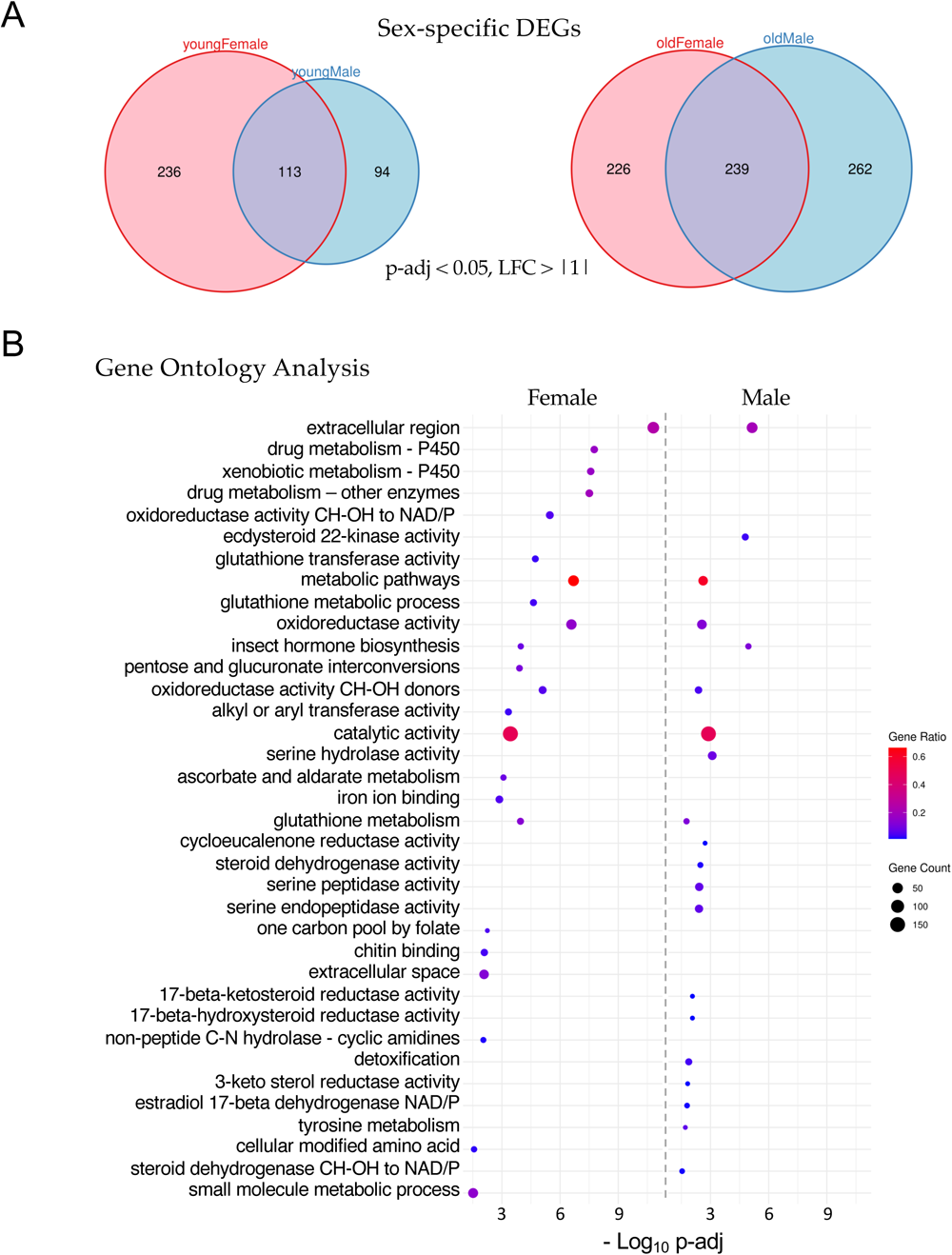
Sex-specific analysis of significant differentially expressed genes. **(A)** Venn diagrams of age-and sex-specific differentially expressed genes, DEGs (p-adj < 0.05 and, LFC > |1|) between *H3.3^K36R^* vs. *H3.3A^null^* fly heads. Left diagram compares DEGs in young female with young male fly heads (0-2 d post-eclosion), and right diagram compares DEGs in old female with old male fly heads (21-25 d). **(B)** Gene Ontology (GO) analysis of the DEGs identified in panel A (old female and old male). The size of each dot is proportional to the number of genes contained within a given ontology term (gene count), and the fraction of those genes scoring significantly (gene ratio) is represented using a heatmap. Adjusted p-values (–Log10 transformed) for each GO term were calculated and plotted separately for both females and males.

### A statistical interaction model highlights defects in innate immune signaling

Though informative, our comparison to the *Chd1* RNA-seq dataset [25] and our original DESeq2 model were unable to take into account the effects of both age and sex. We therefore wanted to design an experiment to interrogate the differential effects that age and sex have on *H3.3^K36R^* flies, relative to *H3.3A^null^* controls. As shown in Fig. 4A, we used DESeq2 and R to create a multifactorial statistical model with two genotypes (*H3.3^K36R^* and *H3.3A^null^*), four conditions (i.e. with age and sex as covariates) and the interaction between age and genotype (age:genotype). This new model places higher weight on gene expression differences that result from the interaction between genotype and condition, rather than by identifying significant differences within a genotype in two separate conditions (for additional details, see Methods). The interaction model thus accounts for the fact that the control genotype could have a different baseline (Fig. 4A) for each of the various conditions (young and old). Strikingly, the age:genotype interaction model reduced the total number of statistically signifcant gene expression changes (p-adj < 0.05) from 3,256 down to 483 transcripts (Fig. 4B and Table S1). Only 312 out of the 1812 ‘Old-only’ DEGs classified in the previous comparison (Fig. 2D) were found to be significant in the interaction model, and 23 new DEGs were uncovered (Fig. 4B).

**Figure 4.**
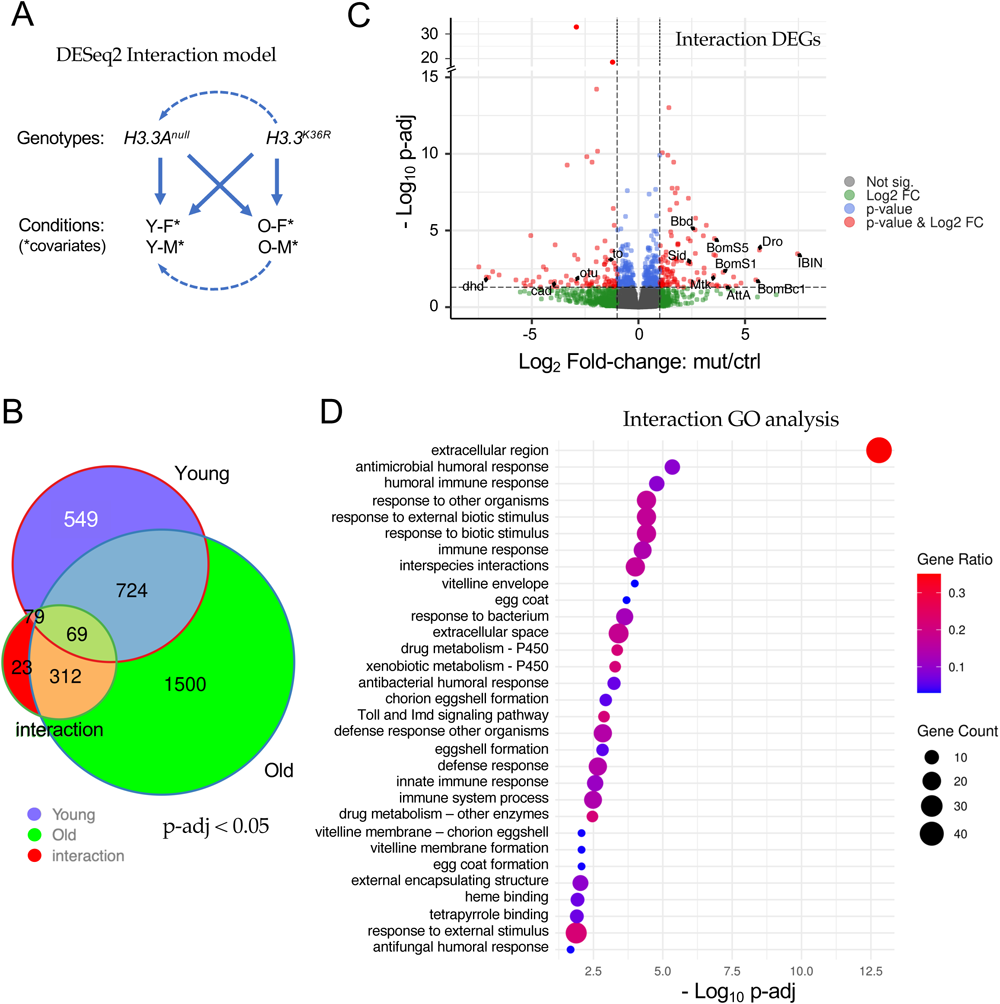
Analysis of the interaction between age and genotype in old and young fly heads. **(A)** Graphical representation of the DESeq2 model including interaction terms (design = sex + age + genotype + age:genotype). Solid arrows represent genotype:condition terms, dotted arrows illustrate how the model sets reference levels. **(B)** Venn diagram of differentially expressed genes (p-adj < 0.05) in young *H3.3^K36R^* vs. *H3.3A^null^* (0-2 d post-eclosion) and old *H3.3^K36R^* vs. *H3.3A^null^* fly heads (21-25 d), compared to those identified using the interaction model (age:genotype). **(C)** Volcano plot of RNA-seq data obtained using the interaction model. Dotted lines represent significance cutoffs for adjusted p-value and log2 fold-change. Dots represent individual genes, color coded according to the key at right. **(D)** Gene Ontology (GO) analysis of DEGs (padj < 0.05 and LFC > |1|) that were determined using the interaction model. Adjusted p-values (–Log10 transformed) for each GO term were calculated and plotted. The size of each dot is proportional to the number of genes contained within a given ontology term (gene count), and the fraction of those genes scoring significantly (gene ratio) is represented using a heatmap.

A volcano plot of all the genes identified using the age:genotype interaction is shown in Fig. 4C. As illustrated, the distribution of the Interaction DEGs (p-adj < 0.05, LFC > |1|, red dots) was roughly equal between up-and down-regulated genes. Notable among the genes whose expression went up are are numerous downstream targets of NF-kB signaling. These include the antimicrobial peptides (AMPs) Drosomycin (Drs), Drosocin (Dro), Attacin A (AttA), Metchnikowin (Mtk) and Bomanins S1, S2, S3 and S5, Bc1, Bc3, T1, T2 T3, (Fig. 4C and Table S1). Other well known immune effectors include IBIN/CG44404, IM4/CG15231, IM14/CG33990, Bbd/CG18067, and SPH93/CG44404. These host defense factors are encoded by well known anti-bacterial and anti-fungal response genes that are targeted by the Toll and IMD signaling pathways [32, 33]. Indeed, the NF-kB ortholog dorsal (dl) is itself among the genes that are differentially upregulated in the mutants (Table S1).

Among the genes that were significantly reduced in expression are several notable transcription factors, including ovarian tumor (otu/CG12743), and caudal (cad/CG1759). Cad and otu are negative regulators of immune (IMD) signaling that, in the adult animal, have been implicated in the maintenance of homeostasis [34–36]. Thus not only are the downstream targets of NF-kB upregulated, but genes involved in dampening or turning off these pathways are also downregulated. Another indicator of a runaway immune response in the brains of aging K36R mutants is highlighted by a 5-fold upregulation of the *Stress-induced DNase* gene, *Sid/CG9989* (Table S1). Sid protein is highly induced by bacterial infection and is thought to protect cells from toxic effects of excess nucleic acids that are released during a vigorous immune challenge [37].

Consistent with the findings outlined above, GO analysis of the Interaction DEGs (using a more stringent p-adj < 0.05 and LFC > |1| cutoff) identified these and many other genes implicating a hyperactivation of immune response pathways in the absence of an external challenge (Fig. 4D). Although these pathways have been clearly implicated in numerous other studies of senescence and aging, the work here provides the first evidence for linking them directly to H3.3K36-dependent processes in animals (see Discussion).

### Age-dependent loss of heterochromatic silencing

Previously, we showed that *H3.3^K36R^* mutants eclose at significantly higher frequencies than those that lack both copies of *H3.3*, strongly suggesting that H3.3K36R protein is incorporated into chromatin and is partially functional. Chromatin profiling of wandering third instar wing discs showed that the *H3.3^K36R^* mutation had little effect on H3K27me3 levels overall, yet the mutant adults displayed mild homeotic transformations [17]. Hence, we were interested in determining which types of chromatin might be affected by loss of H3.3 lysine-36.

Kharchenko and colleagues have classified nine different *Drosophila* chromatin “states” that are based on a wide variety of underlying histone marks and binding factors [38]. We used genome coordinates, derived from profiling data in BG3 larval brain cells, that have been established for each of these states to carry out an analysis of the genes that were differentially expressed in our K36R mutants. Rather than focusing on a small subset of DEGs whose contribution to the phenotype is uncertain, we broadened this analysis to the union set of young and old DEGs (Fig. 2D). Genes whose coordinates did not overlap with those of any single chromatin state by at least 50% were binned into a tenth (NA) category.

As shown in Fig. 5A, we plotted the mean Log2 fold-change over time for the DEGs that fell into each of these ten categories (Chromatin States). The plot on the left within each color grouping measures fold-change between young K36R mutants vs controls, whereas the plot on the right does the same for the older animals. The color code and brief descriptions of the characteristics for each chromatin state [38] are provided in Fig. 5B. As illustrated, Chromatin States 1-3, which comprise roughly 30% of the genome and are characterized by ‘active’ marks, are relatively unchanged from young to old animals (Fig. 5A). DEGs in States 4, 7 and 8 are characterized by less active marks and showed the greatest degree of change over adult lifespan. In contrast, DEGs in States 5 and 6 trended towards lower expression levels over time (Fig. 5A). State 5 is reflective of the dosage-compensated male X chromosome (characterized primarily by high levels of H4K16ac), and the K36R DEGs in this category were highly enriched (245/266 genes) on the X chromsome, Fig. 5C. Base mean expression levels for DEGs in each state are provided for purposes of comparison (Fig. S4A). Surprisingly, the downward trend in expression levels in DEGs from State 5 was apparently not driven by males (Fig. S4B), suggesting that H3.3K36 may not play a direct role in X chromosome dosage compensation.

**Figure 5.**
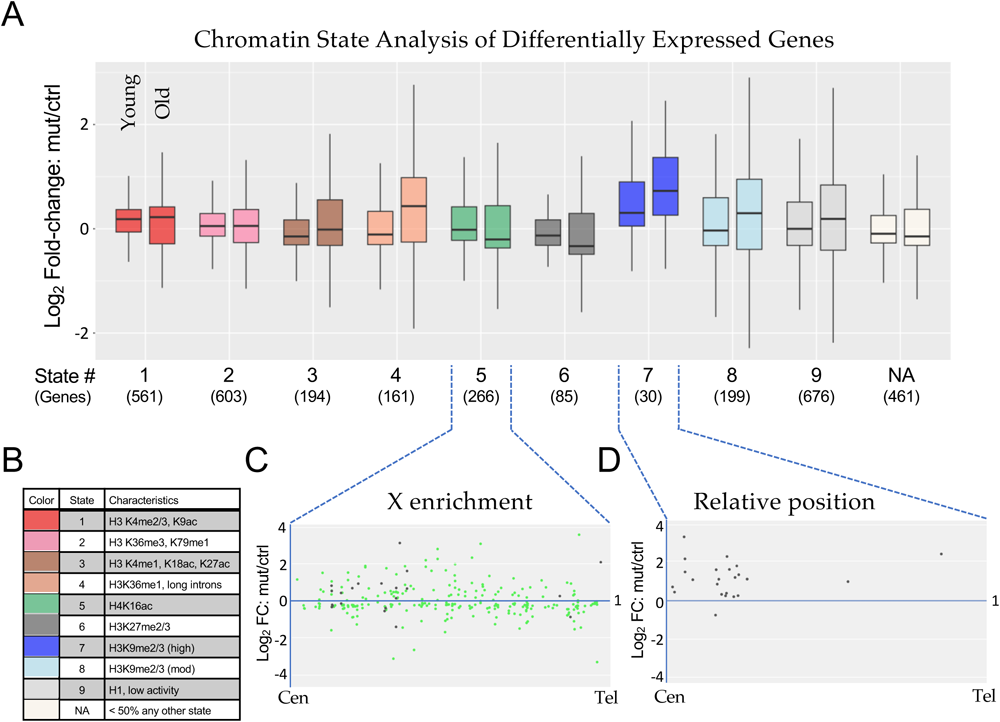
Chromatin state analysis of differentially expressed genes. **(A)** The combined set of differentially expressed genes (adjusted p-value < 0.05) from the non-interaction DESeq2 model for both Young and Old conditions for *H3.3^K36R^* mutants relative to *H3.3A^null^* controls was binned by predominant chromatin state [38]. Genes were binned to a given state if > 50% of the gene was marked by that state. The “NA” category was comprised of genes where no state color was > 50% of gene length (406 genes) or if chromatin states were undetermined when the original chromatin state study was published (55 genes). The mut/ctrl Log2 fold-change value for binned genes was plotted separately for Young and Old conditions. **(B)** A legend for colors corresponding to each chromatin state depicted in panel A, with representative histone marks for each state. **(C** and **D)** For genes in States 5 and 7, mut/ctrl Log2 fold-change values for the Old condition were plotted versus relative position along chromosome arms for all chromosomes, from centromere (Cen) to telomere (Tel). For State 5, the X chromosome genes are colored green, and autosomal genes, black.

The DEGs in State 7 showed the greatest degree of expression change in young adults as well as in change over time (Fig. 5A). State 7 is characterized by high levels of heterochromatic H3K9me2/3 marks (Fig. 5B). Although there was no single chromosome arm that harbored the bulk of these genes, 7/30 DEGs were located on the primarily heterochromatic 4^th^ chromosome. Interestingly, 21 of the remaining 23 DEGs were located in centromere-proximal locations along their respective chromosome arms. A metaplot of these 23 genes from State 7 is shown in Fig. 5D, showing that all but one of them were derepressed in the mutant. Note that the genome build for these regions is incomplete, and so the relative position of the centromere is not the same for each arm. Thus the metaplot is not meant to depict the precise location of each DEG, but rather to illustrate its general positioning along the arm. Nevertheless, the results collectively provide strong evidence that loss of H3.3 K36 results in derepression of genes located in heterochromatic regions of the genome.

### *H3.3^K36R^* mutants exhibit position effects in the adult eye

In contrast with its typical role as a marker of active genes along chromosome arms, H3.3 is also known to be deposited by the ATRX•Daxx complex to maintain repressive pericentric and (sub)telomeric chromatin [39–41]. In mouse embryonic stem cells, loss of *H3f3*a and *H3f3b* (orthologs of *H3.3A* and *H3.3B* in flies) results in reduced levels of H3K9me3 and ATRX at telomeres [42]. The chromatin state analysis of *H3.3^K36R^* DEGs shown in Fig. 5 clearly implicates a disruption to H3K9-mediated silencing, particularly in centromere-proximal regions. However, due to the paucity of protein coding genes located near telomeres, this approach is unable to evaluate the transcriptional status of telomere-proximal heterochromatin.

The fruitfly genome is rare among eukaryotes because it does not encode a telomerase RNP [43]. Instead, telomere length is maintained by controlled retrotransposition of *HeT-A (Healing Transposon A)*, *TART (Telomere Associated Retrotransposon)* and *TAHRE (Telomere Associated and HeT-A Related)* elements into a telomere-proximal region (Fig. 6A). Telomeric chromatin is comprised of two distinct epigenetic compartments that are believed to perform different functions. The so-called HTT (*HeT-A, TART, TAHRE*) array (Fig. 6A), is primarily marked by H3K9 methylation that is thought to be required for proper expression of the HTT array [43, 44]. Adjacent to the array are the Telomere Associated Sequences (TAS, Fig. 6A), which contain 40-60 copies of telomere-specific satellite repeats per chromosome arm [43, 45]. Interestingly, TAS repeats are marked by repressive H3K27me3 and Polycomb Group (PcG) proteins [46].

**Figure 6.**
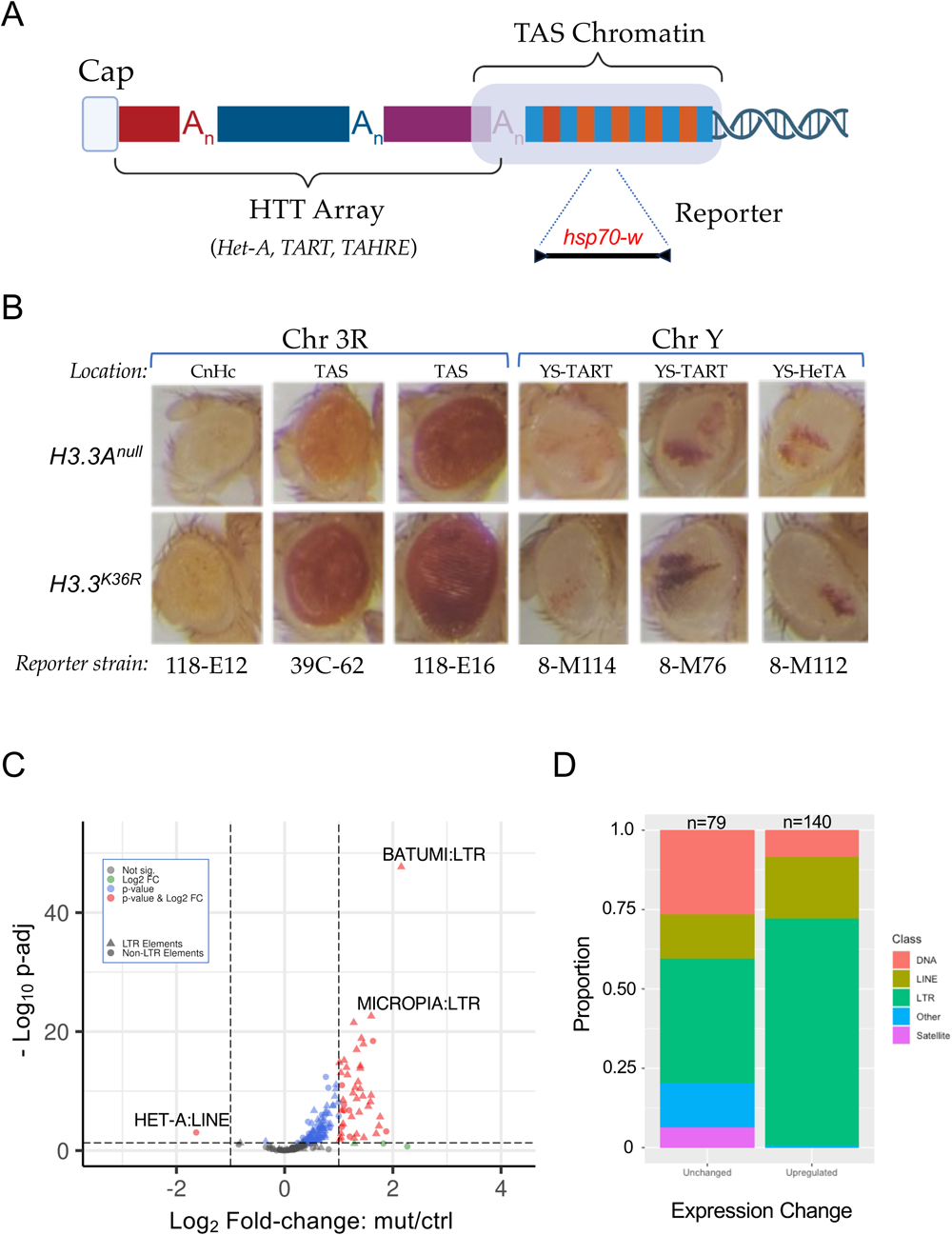
Telomeric position effect and transposable element expression analysis. **(A)** Cartoon depicting key features of *Drosophila* telomeres. At the distal end of the telomere, a capping complex assembles adjacent to arrays of *Het-A, TAHRE,* and *TART* retrotransposons (HTT array). Proximal to that is the TAS (Telomere Associated Sequence) Chromatin, which is characterized by the presence of complex satellite repeats that vary by chromosome. **(B)** Images of adult eyes. For each column, representative examples are shown for a particular reporter strain. Rows correspond to genotype (*H3.3A^null^* control or *H3.3^K36R^* mutant). The location and type of chromatin for each reporter strain is also indicated: Pericentric heterochromatin (CnHc), Telomere Associated Sequences (TAS), YS-TART (Y chromosome TART), and YS-HeTA (Y chromosome HeT-A). n-values for each experiment ranged from 16 to 44, see Fig. S5 for details. **(C)** Volcano plot of annotated transposable elements (TEs) analyzed using the TEcounts tool within TEtranscripts. Each dot or triangle represents the combined reads from all loci for a given TE subtype. Dotted lines represent significance cutoffs for adjusted p-value and Log_2_ fold-change. Symbols are color coded according to the key in the upper left. LTR-type transposons are represented by triangles, non-LTR elements are shown as dots. **(D)** Stacked bar graph, binned by whether TEs were upregulated or unchanged in the *H3.3^K36R^* mutant. The relative proportion of a given type of transposable element (DNA, LINE, LTR, Satellite, and Other) in each bin are shaded as indicated in the legend.

In order to assay the functional consequences of *H3.3^K36R^* mutation at telomeres, we utilized a miniwhite reporter system pioneered by Wallrath and Elgin [47]. These and other reporters have been widely used to analyze position effect variegation (PEV) of the *white* gene in the adult *Drosophila* eye (Fig. 6A). We ingressed six different *hsp70-w* reporter transgenes, located either on 3R or YS, into the background of *H3.3A^null^* or *H3.3^K36R^* mutants. Five of these lines contain insertions in telomeric chromatin (two in TAS on 3R; two in the HTT array of the Y chromosome), and one of them (118-E12) is inserted near centric heterochromatin on 3R (Fig. 6B). Surprisingly, none of the HTT reporters on the Y chromosome showed appreciable differences in the intensity or extent of PEV in the K36R mutant eyes, compared to the controls (Fig. 6B). In contrast, each of the TAS reporters on 3R showed an increase in the overall intensity of red eye pigmentation compared to the basal levels of the control animals. Also note that Y chromosome reporters showed significant variegation, whereas the 3R lines had more uniform expression levels. Although the basal level of pigmentation in 118-E6 is quite high, and that of line 118-E12 is rather low, we consistently found slightly higher levels in the K36R animals (Fig. 6B and Fig. S4). This effect is more clearly seen in line 39C-62, which has moderate basal levels and demonstrably higher levels in the presence of the K36R mutation (Fig. 6B). These observations indicate that H3.3K36 may play a role in maintaining a suppressive chromatin environment within TAS as well as centric heterochromatin.

### Loss of H3.3 K36 reduces expression of telomere-specific retroposons, but causes derepression of many other transposable elements

The role of ATRX•Daxx-mediated H3.3 deposition in promoting and maintaining the silencing of transposable elements (TEs) is a well documented feature of higher eukaryotic genomes [48–50]. Thus we were curious to directly determine whether transcription of telomeric transposons was altered. Because transposons are repetitive elements, traditional RNA-seq pipelines are not set up to properly apportion short sequencing reads to repeat-rich regions of the genome. Hence, differential analysis of transposable element (TE) expression was performed using the open source software package, TEtranscripts [51]. A volcano plot of these data is shown in Fig. 6C, revealing a dramatic TE derepression phenotype. Prominent among the classes of *upregulated* TEs are LTR-type retrotransposons (Figs. 6C,D). Increasing mobilization of transposons leading to DNA damage has also been associated with aging [52, 53]. These data suggest that activation of repressed TEs may contribute to the reduced lifespan of *H3.3^K36R^* mutant animals.

Conspicuous among the TE families that are *downregulated* in K36R mutants is the *HeT-A* family of LINE-like (non-LTR) retroposons involved in telomere length maintenance. As shown in Fig. 6C, expression of transcripts corresponding to the *HeT-A* family is dramatically reduced in old K36R mutant vs. control fly heads. Note that we also found that expression of both *HeT-A* and *TART* elements was significantly reduced in the old K36R mutants if we employed the more commonly-used featureCounts tool instead of TEcounts (see Tables S2 and S3). Among the three telomeric retrotransposons, *HeT-A* elements are particularly important because the *HeT-A* Orf1p/Gag protein recruits its *TART* counterpart to chromosome ends [54, 55]. Thus, reduced expression of *HeT-A* elements would be expected to have serious consequences for telomeric stability [56]. Telomere shortening is a well known hallmark of cellular senescence; these data are consistent with the notion that defects in telomere maintenance contribute to the observed reduced lifespan phenotype in *H3.3^K36R^* animals.

### Hyperacetlyation of histone H4 in H3.3K36R chromatin

Our finding of impaired heterochromatic silencing in H3.3K36R mutants aligns well with previous observations of senescence-associated heterochromatin defects in aging insects [57, 58] as well as in senescent mammalian cells [13, 59]. Another hallmark of senescent cells and tissues are changes in the acetylation status of histone H4 (reviewed in [60–62]). Indeed, these changes in histone acetylation patterns are often already apparent when animals reach midlife [63]. Using a multi-gene replacement strategy to analyze functions of the replication-dependent H3.2 histones [64, 65], we previously showed that an H3.2K36R mutation caused global hyperacetylation of H4 [66]. To determine if an H3.3K36R substitution might have a similar effect, we measured overall histone H4 acetylation levels in mutant and control animals by western blotting.

As shown in Figure 7, we assayed bulk levels of histone H4ac and H3 (loading control) and found that acetylated H4 was significantly higher in H3.3K36R mutants compared to H3.3Anull or yw control animals. Confirming the previously identified link between H3K36me and H4ac in yeast, these findings are also consistent with the idea that Set2-mediated K36 trimethylation recruits the histone deacetylase complex, RPD3S [67, 68]. Histone hyperacetylation has long been regarded as a dynamic characteristic of transcriptionally active genes, a property associated with elongating polymerases [69].

**Figure 7.**
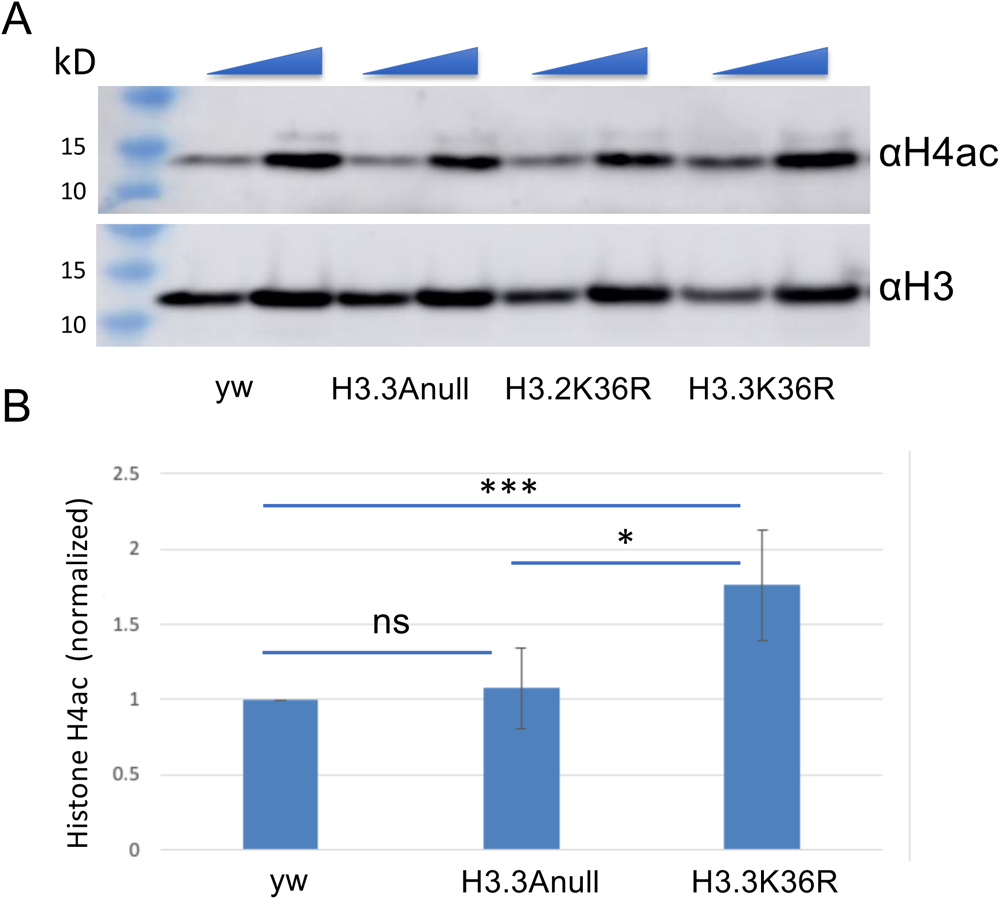
Hyperacetlyation of histone H4 in H3.3K36R mutants. Western blotting for pan H4 acetylation (H4ac) and anti-H3 on lysates from WL3 larvae. (**A**) Representative gel image. yw and H3.3Anull controls are shown along with H3.3K36R mutants. The H3.2K36R lanes are provided for visual comparison but are not quantified (see Methods for details). As indicated by the triangles at the top edge for each pair of lanes per genotype, either 20 or 40 ug of total protein per lane was loaded. (**B**) Quantification of the data from three independent replicates. Mean pixel intensity for each band was measured, and ratio of H4ac/H3 was computed per replicate. For each experiment, the H4ac/H3 ratio of mutant genotypes was normalized to the *H3.3A^null^* control. For each genotype, normalized means and SD were plotted. Significance between raw H3K27me3/H3 ratios was assessed by paired one-way ANOVA, followed by Friedman tests comparing mutant genotypes to *H3.3A^null^* controls. *P* values are indicated as follows: *** < 0.001; * < 0.05; ns = not significant.

Recently, an investigation of diverse metazoan species, including *Drosophila*, found that transcription elongation rates naturally increase with organismal age, and that reducing Pol II speed extends lifespan [70]. Given that histone H4 acetylation is known to destabilize nucleosomes during transcription (reviewed in [71]), activities that serve to hyperacetylate histones would be expected to increase polymerase elongation rates. Taken together, our findings are consistent with the notion that hyperacetylation of H4 may contribute to the premature lethality of H3.3K36R mutant flies.

## Discussion

Aging is a complex process that involves multiple factors contributing to the decline of cellular and organismal functions, ultimately leading to decreased lifespan. Cellular senescence is characterized by numerous features, including irreversible cell cycle arrest, altered gene expression patterns, metabolic changes, and the acquisition of a senescence-associated secretory phenotype (SASP). Senescent cells secrete pro-inflammatory cytokines, growth factors, and matrix remodeling enzymes, contributing to chronic inflammation, tissue dysfunction, and age-related pathologies. Epigenetic changes, including histone modifications, have been identified as crucial contributors to aging-related processes. In this study, we focused on the lysine-36 residue of the replication-independent histone protein, H3.3. Our results demonstrate that expression of H3.3K36R as the sole source of H3.3 protein causes a significant reduction in adult lifespan and dysregulation of the genomic and transcriptomic architecture. Genetic rescue of the longevity defect upon transgenic co-expression of wild-type *H3.3B* suggests a direct role of lysine-36 in supporting longevity. Additionally, our transcriptome profiling of young and aged adult fly heads reveals age-dependent misexpression of innate immune target genes, hyperacetlyation of histone H4 and derepression of heterochromatic transcripts in H3.3K36R mutants.

### Role of Toll/Imd Pathways in Aging and Immuno-senescence

The Toll/Imd pathways are thought to play a vital role in regulating immunity and are intertwined with the aging process. Knockdown of Toll/Imd pathway members extends lifespan, whereas their overexpression shortens it [72]. Immuno-senescence, characterized by age-dependent decline in immune function, is a hallmark of the aging process [73, 74]. Interestingly, both sexes become more susceptible to infection as they age, with different underlying mechanisms. For males, susceptibility is primarily posited to be caused by deterioration in barrier defenses, whereas for females, systemic immune defense senescence is thought to be the major contributor [75].

Our findings are also consistent with observations in mammals, wherein NF-κB activity increases over time in the brains of middle-aged and older mice relative to young mice [76]. The upregulation of AMP expression in aged *Drosophila* [77] further supports the connection between immune signaling and the aging process. Notably, downregulation of NF-κB has been shown to mobilize nutrients and extend lifespan, whereas its overexpression has been linked to neurodegeneration [76, 77]. Our investigation into the molecular consequences of the *H3.3^K36R^* substitution highlights its involvement in regulating critical aspects of immune signaling and heterochromatic silencing. Using a statistical interaction model, we discerned the effects of age and sex on gene expression in *H3.3^K36R^* mutants compared to controls. Notably, the expression of downstream targets of NF-κB signaling, such as antimicrobial peptides and other immune effectors, were upregulated in the mutants. This suggests that H3.3K36 may play a role in suppressing immune responses in the absence of external immune challenge, thereby contributing to the maintenance of cellular homeostasis in the adult brain.

### H3.3K36, Retrotransposition and Senescence

Understanding the intricate interplay between chromatin dynamics and the aging process should offer valuable insights into mechanisms of aging and age-related diseases. Upon loss of H3.3K36, our analysis has revealed a disruption of H3K9-mediated silencing in both centromere-and telomere-proximal regions. This includes a robust downregulation of non-LTR retrotransposons that are critical for maintaining telomere length [54, 55]. In contrast, the observed derepression of genes and LTR-type retroposons in heterochromatic regions provides strong evidence that H3.3K36 is involved in maintaining repressive pericentric and subtelomeric chromatin, possibly through ATRX•Daxx-mediated deposition. Consistent with these observations, Groth and colleages have recently shown that DAXX provides additional functionality to the histone chaperone system by recruiting H3K9 methyltransferases [78]. Even more recently, the same group showed that a failure to re-establish H3K9me in post-replicative mammalian chromatin causes derepression of LTR-type retroposons [79]. Together, the data suggest that, in certain parts of the genome, H3.3K36 plays a conserved and specific role in silencing these so-called genomic parasites.

We also observed that mutation of H3.3K36 results in reduced repression of reporters in H3K27me3 marked TAS, and reduced expression of chromatin State 6 genes (marked by H3K27me3 and Pc proteins). These findings are in line with our recent work suggesting a role for H3.3K36 in supporting the function of genes marked by H3K27me3 [17]. Studies in many organisms point to gene regulation by PcG proteins as another pathway that influences longevity [14, 80–82]. However, the evidence is inconsistent between organisms, and much remains to be learned mechanistically. Further work clarifying the relationship between H3.3K36, Polycomb mediated gene repression, and aging would be an interesting area of future investigation.

It is intriguing to speculate on mechanisms through which H3.3K36 might support retrotransposon expression at telomeres. One possibility is that this residue facilitates expression of piRNA pathway genes. The piRNA pathway functions to suppress transposon activity in the germline, including activity of the HTT array [83–86]. Consistent with our observation of downregulation of *HeT-A* and widespread derepression of other TE families in *H3.3^K36R^* mutants, previous studies have observed differences in regulation of telomeric piRNA clusters compared to those that target non-telomeric TEs [84, 85]. Furthermore, analysis of a dominant negative *H3.3^K36M^* model (causing reduction in methylation of both H3.2K36 and H3.3K36), showed that telomeric retrotransposons were downregulated by piRNA pathway genes [87]. In *H3.3^K36M^* mutant eye-antennal discs, suppression of *HeT-A* levels was elicited by upregulation of the piRNA promoting gene, *krimp*. Importantly, the suppression of *HeT-A* levels in these mutants could be rescued solely by reducing *krimp* expression [87]. Of note, we also observe upregulation of *krimp* in old *H3.3^K36R^* mutants (log2 fold change = ∼1.3), as well as another piRNA pathway gene, *Ago3* (log2 fold change = ∼1.2), see Table S1 for details.

Another mechanism through which H3.3K36 might sustain expression of the HTT array is by promoting Jil-1 kinase function. Phosphorylation of the Jil-1 target, histone H3 serine 10, antagonizes both methylation of H3K9 and gene repression [88, 89]. Jil-1 is a positive regulator of HTT transcript levels [56, 90]. Moreover, *Jil-1* and *H3.3^K36R^* mutants both exhibit repression of *HeT-A* transcription and suppression of TAS-reporter transgenes ([56]; this work). Recently, an H3K36me3 reader protein, Jasper, has emerged as an important factor for stabilization and recruitment of Jil-1 [91, 92]. *Jasper* mutants exhibit both an increase in H3K9me2 and a decrease in expression of *HeT-A* and *TART* transposons [91]. However, unlike *H3.3^K36R^* mutants, *Jil-1* and *Jasper* mutants do not exhibit widespread derepression of transposons in other parts of the genome [91, 92]. An interesting area of future study would be to determine whether this difference might be due to activities specific to H3.2K36 versus H3.3K36.

In summary, we find that lysine-36 of *Drosophila* histone H3.3 is crucial for supporting adult longevity by regulating immune signaling, heterochromatic silencing, and retrotransposon repression. The dysregulation of these processes in H3.3K36R mutants likely contributes to age-related changes in gene expression and chromatin organization, ultimately impacting organismal lifespan. Further investigations into the mechanistic details of how H3.3K36 functions in these pathways will shed light on the broader role of histone modifications in the aging process. Additionally, exploring the conservation of these mechanisms in other organisms, including mammals, may have implications for understanding aging and age-related diseases in humans. Future studies of this process in model organisms may also provide potential therapeutic targets to mitigate age-related decline and improve healthy aging.

## Materials and Methods

### Fly Husbandry and Genetics

We generated *H3.3^K36R^* mutants through creation of a K36R missense mutation at *H3.3B*, and combining it with an *H3.3A* deletion (*H3.3AΔ*). Both *H3.3A^null^* and *H3.3^K36R^* animals are obtained through a cross and negative selection for CyO, twGFP chromosome. *H3.3A^null^* animals are made by crossing yw; H3.3^2x1^/CyO, twGFP with yw; Df(2L)Bsc110/CyO, twGFP. *H3.3^K36R^* animals are made by crossing yw, H3.3B^K36R^; H3.3^2x1^/cyO, twGFP with yw, H3.3B^K36R^; Df(2L)Bsc110/CyO, twGFP. The H3.3A^2x1^ and Df(2L)Bsc110 alleles were obtained from Bloomington Drosophila Stock Center (#68240, #8835), whereas the *H3.3B^K36R^* mutation was generated as described in [17]. Crosses were carried out in cages sealed with grape juice agar plates with supplemental yeast paste. Plates were replaced daily and GFP-negative larvae were sorted into vials of cornmeal-molasses food. Experimental flies were raised at 25°C, and all experiments were performed at 25°C unless otherwise noted.

### Adult lifespan assays

To assess longevity, parental stocks were mated on grape juice agar plates and larvae were sorted into vials of molasses food. For each replicate, 50 larvae were sorted into fresh food vials (total of 4-6 replicates per genotype) and the fraction (%) of animals that pupated and eclosed was calculated. The mean and standard deviation for each genotype was plotted with GraphPad GraphPad Prism software. Following eclosion, male and female flies were kept in separate vials. Animals that eclosed within 48hrs of each other were considered the same age. For the adult lifespan assays, each vial contained twenty or fewer flies to avoid crowding, and was incubated on its side to reduce accidental deaths from animals getting stuck in the food. Dead flies were removed and survivors flipped into fresh food every 2-3 days to prevent death due to poor conditions. Survivors were re-counted and recorded after each flip. Flies that were killed or escaped during the experiment were removed from the count.

### Fly head RNA extraction and RNA Sequencing

*H3.3^K36R^* and *H3.3^null^* animals were generated as above and were separated into groups based on the date of eclosion. Flies that eclosed within 48 hrs of each other were considered an age cohort. Young cohorts (0-2 days post-eclosion) were processed without further aging, whereas old cohorts were processed after three weeks (21-25 days post-eclosion). Heads were removed from the rest of the carcass and total RNA was extracted. For each bioligical replicate, 10-25 flies were anesthetized and placed in a 15 mL tube. The tube was submerged in liquid nitrogen for 5-10 seconds and then vortexed for 5-10 seconds to separate heads from carcasses. This process was repeated for a total of 5 times before pouring the contents of the tube into a set of stacked sieves (Cole-Parmer, 3“-Diameter Sieve, Full Height Stainless Steel, No. 25 on top of Cole-Parmer, 3”-Diameter Sieve, Full Height Stainless Steel, No. 40). Fly heads were recovered from the bottom sieve and either stored at −80°C or immediately used for RNA extraction. Fly heads were crushed in TRIzol reagent (Thermo Fisher) and RNA was purified with the RNA clean & concentrator-5 kit (Zymo research), and equivalent amounts of RNA per replicate were used for cDNA library construction. Total (Ribo-minus) RNA-seq libraries, six (3 male, 3 female) replicates per genotype, were prepared using a Universal RNA-seq library kit with NuQuant (Tecan). Sequencing was performed on a NovaSeq 6000SP (Illumina) producing 100 nt, paired-end reads.

### RNA-seq Analysis and Data Availability

Raw and processed RNA-seq data were deposited to Gene Expression Omnibus (GEO) under accession GSE244389. Quality control was performed with FastQC, ver. fastqc/0.11.7 [93], and reads were aligned using STAR [94] to genome build dm6. Aligned reads were counted with featureCounts [95] and differential gene expression analysis was performed with DESeq2 (v1.34.0) [96]. One outlying sample was removed from the downstream analysis. Standard DESeq2 protocols were used to determine statistical significance of differential gene expression in Fig. 2 and 3. The statistical model used in Fig. 4 (design = sex + age + genotype + age:genotype) takes into account changes in the effect size and direction between the two conditions and the genotypes. Thus this so-called ‘interaction model’ places greater statistical weight on gene expression differences that result from the interaction between genotype and condition.

Gene ontology analyses were performed using g:Profiler [97]. Differential transposon analysis was performed using TEtranscripts [51]. The statistical model used in this case included sex and genotype, as we chose to compare old mutants to old control animals.

### Assignment of Differentially Expressed Genes to Chromatin States

Files containing Chromatin State information for larval BG3 cells, as defined previously [38], were downloaded from the modENCODE website. Gene interval coordinates for transcription-start and −end sites (TSSs and TESs) were intersected with the Chromatin State files using BEDTools [98] to assign genes to a given state if they were covered by greater than 50% over its length. Those wherein no State exceeded 50% coverage were labeled as mixed. Genes were binned by state and plotted by Log2 fold change values determined by DESeq2 [96] using the non-interaction model.

### Position Effect Variegation Experiments

Position effect variegation (PEV) reporter lines were obtained from the Bloomington Drosophila Stock Center: 118E-12, stock #84015; 39C-62, #44261; 118E-16, #84106; 8-M114, #84095; 8-M76, #84096; and 8-M112, #84093. These PEV lines carry a variegating hsp70::mini-white insertion [47]. To assess the role of H3.3K36 in heterochromatin spreading, PEV males with an H3.3A^2x1^/CyO 2^nd^ chromosome were crossed to yw, H3.3BK36R; Df(Bsc110)/CyO, twGFP (H3.3BK36R) or yw; Df(Bsc110)/CyO, twGFP (H3.3AΔ) virgin females. Straight-winged male flies were age-matched, and flies that eclosed within 96h of each other were grouped together. Images were taken on the following day, so the flies were 1-5 days post-eclosion.

### Western Blotting

Protein lysates from whole wandering third instar (WL3) larvae (ten animals per replicate) were obtained by homogenization with a micropestle in SUTEB buffer [1% SDS, 8 M urea, 10 mM EDTA (pH 8.0), and 5% β-mercaptoethanol, with 1:20 Halt protease inhibitor cocktail (Thermo Fisher Scientific, no. 78429)]. Chromatin was further disrupted by sonication with a Bioruptor Pico (Diagenode)..

Samples were electrophoresed on 4 to 15% Mini-PROTEAN TGX Stain-Free protein gels (Bio-Rad, #4568084) for 60 min at 100 V. Western blotting was performed using the Bio-Rad Trans Blot Turbo transfer system using the provided buffer (Bio-Rad, no. 10026938) onto a nitrocellulose membrane at 1.3A/25 V for 7 min. Membranes were blocked at RT in 5% milk in TBS-T (Tris buffered saline, Tween 20). Primary antibody incubation was overnight at 4°C in TBS-T with 5% milk with one of the following antibodies: polyclonal rabbit anti-H4 pan acetyl (1:1000; Activemotif, no. 39925) and polyclonal rabbit anti-H3 (Abcam, #1791). For each primary antibody, an anti-rabbit secondary (Sigma-Aldrich, no. 12-348) was used at 1:5000.

Blots were incubated with chemiluminescent detection reagent (Amersham ECL Prime Western Blotting Detection Reagents, GE Healthcare, no. RPN2236) and imaged on an Amersham Imager 600 (GE Healthcare). Between primary antibody antibodies, blots were stripped, rinsed in TBS-T, and incubated with detection reagent to verify removal of antibody before reprobing. Relative band intensity ratios were calculated using NIH ImageJ. Briefly, a box of equal size and dimension was drawn around each band, and integrated density (IntDen) inside the box was recorded. For each blot, a ratio of H4ac/H3 IntDen was calculated per sample lane. In addition, a normalized value for each mutant genotype was calculated by dividing the mutant ratio by that of the control *H3.3A^null^* genotype. Lysates from yellow white (yw) wild-type control animals, along with 12xH3.2K36R mutants. Quantification of the data from 12xH3.2K36R mutants was not used in the quantification due to differences in overall histone gene number (eacho of the other genotypes contain 200x copies of the replication dependent histone genes).

