## Supplemental figures S1-S5 for "Lysine-36 of *Drosophila* histone H3.3 supports adult longevity"

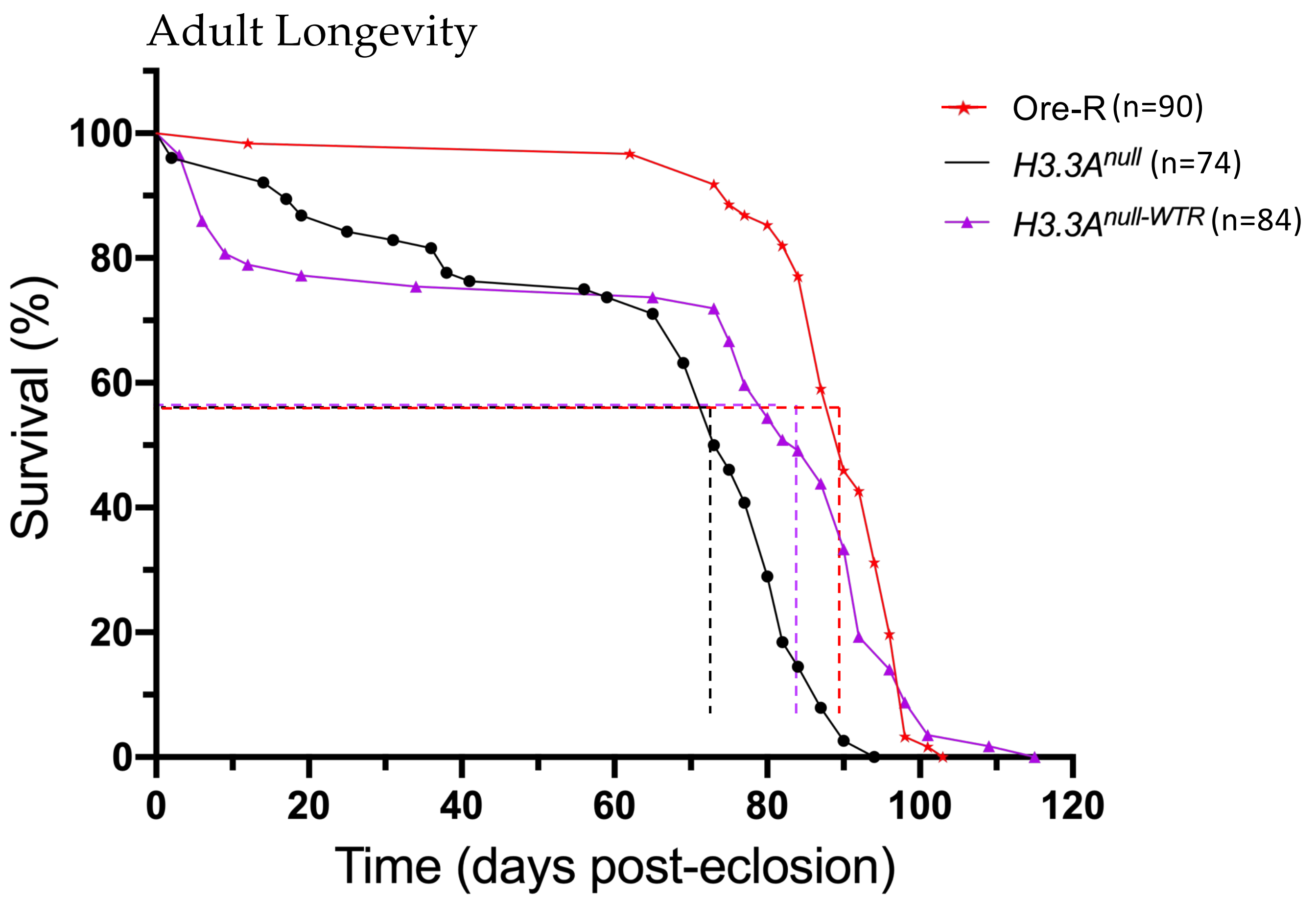


| | Ore-R | *H3.3A^null^* | *H3.3A^null-WTR^* |
|---|---|---|---|
| Ore-R | --- | $< 10^{-6}$ **** | $< 4.3 \times 10^{-3}$ * |
| *H3.3A^null^* | $< 10^{-6}$ **** | --- | $< 4.2 \times 10^{-3}$ * |
| *H3.3A^null-WTR^* | $< 4.3 \times 10^{-3}$ * | $< 4.2 \times 10^{-3}$ * | --- |

**Figure S1.** ***H3.3A$^{null}$*** **control animals are slightly less fit than Oregon-R wild type control animals**. Adult longevity assays for Oregon-R, *H3.3A$^{null}$* and *H3.3A$^{null\text{-}WTR}$* flies. Median lifespan was determined (dotted lines) by identifying the day at which 50% of the animals survived. Statistical comparison of survival curves using Gehan-Breslow-Wilcoxon tests are presented in the accompanying table. A Bonferroni correction for multiple comparisons was employed, resulting in the following adjusted significance values: * $p<0.0167$, ** $p<0.0034$, *** $p<3.4 \times 10^{-4}$, ****$p<3.4 \times 10^{-5}$.

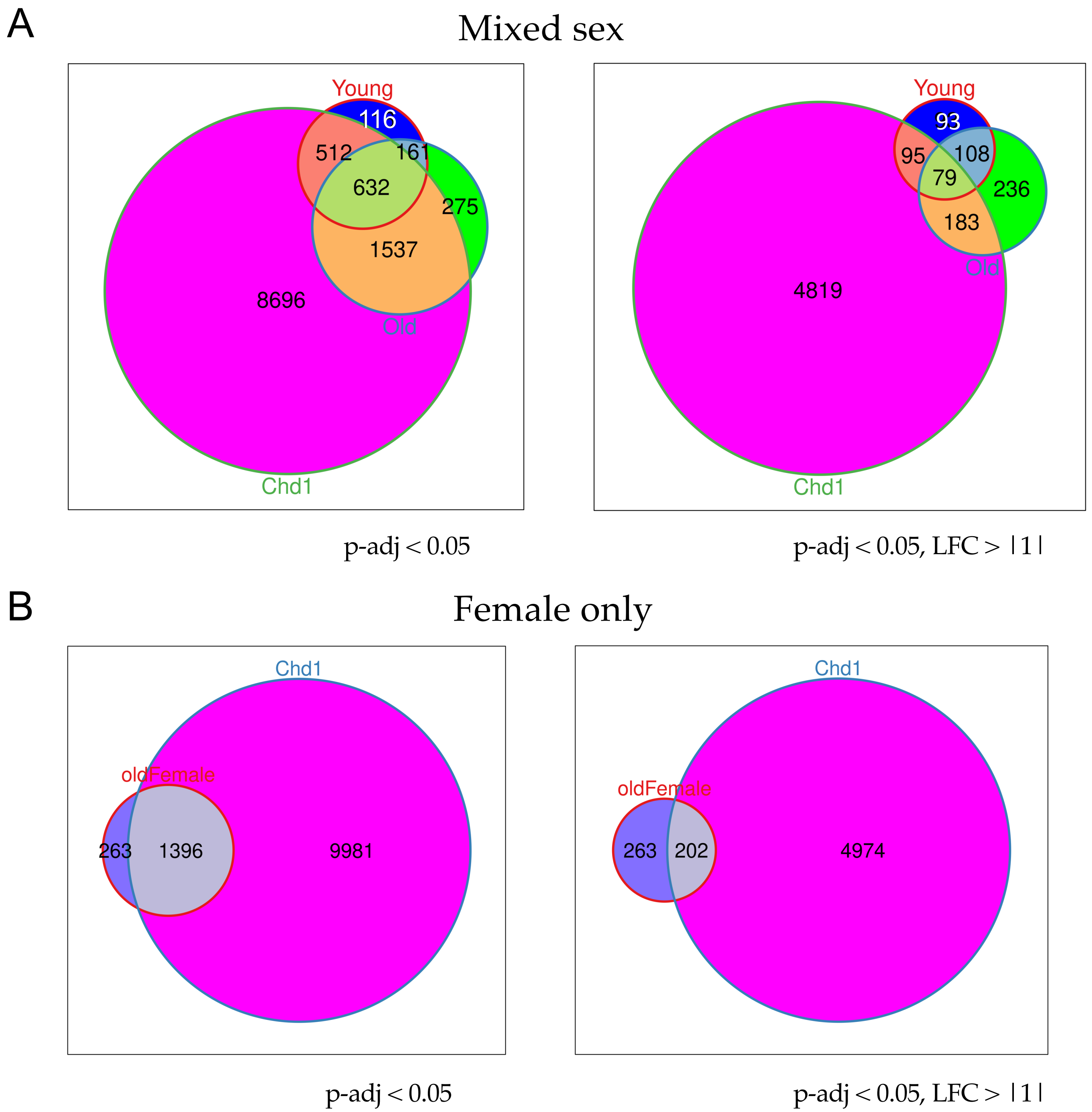


**Figure S2. Overlap of DEGs between $H3.3^{K36R}$ and *Chd1* mutants.** A) Venn diagrams of Young $H3.3^{K36R}/H3.3A^{null}$, Old $H3.3^{K36R}/H3.3A^{null}$, and *Chd1* / Control DEGs from our Mixed Sex (non-interaction) DESeq2 model (Figure 2) and the *Chd1* data reanalyzed with the same bioinformatic workflow. At left, genes with p-adj < 0.05; at right, with an additional cutoff with LFC > |1|. B) Same as in A, except we performed DESeq2 analysis with sex as a variable, displaying DEGs from females only to more closely match the samples in the *Chd1* study.

Figure S2

### Adult Longevity

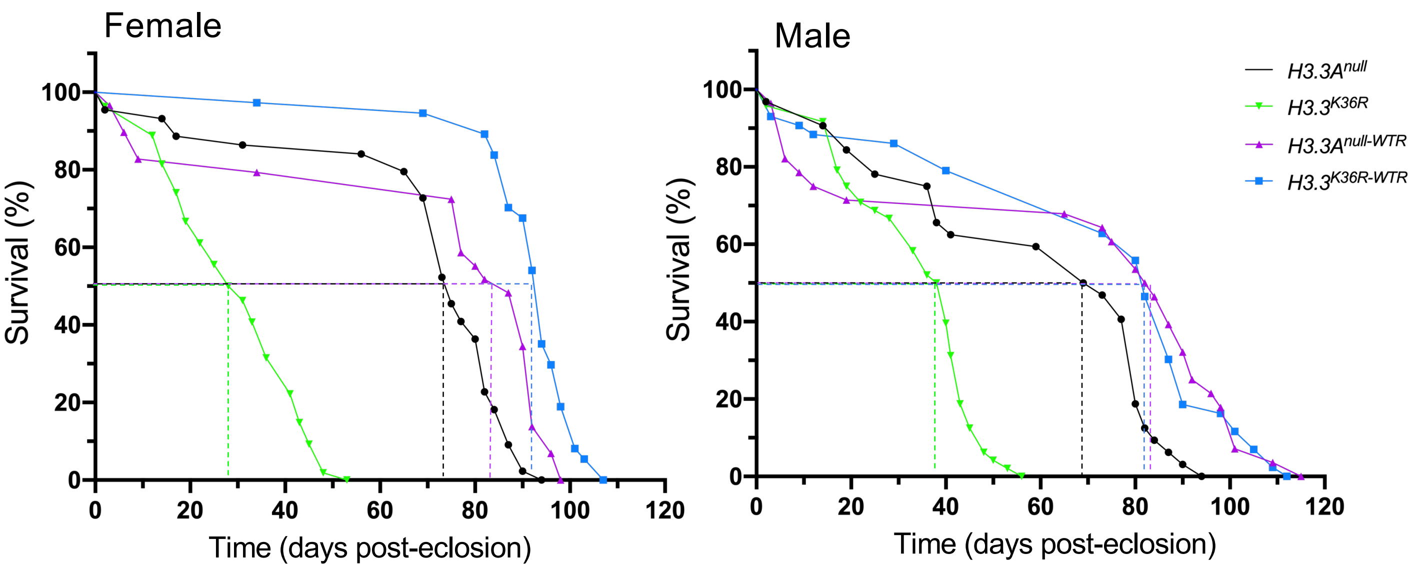


| **Female** | *$H3.3A^{null}$* | *$H3.3^{K36R}$* | *$H3.3A^{null\text{-}WTR}$* | *$H3.3^{K36R\text{-}WTR}$* |
|---|---|---|---|---|
| *$H3.3A^{null}$* | --- | $< 10^{-6}$ **** | ns | --- |
| *$H3.3^{K36R}$* | $< 10^{-6}$ **** | --- | --- | $< 10^{-6}$ **** |
| *$H3.3A^{null\text{-}WTR}$* | ns | --- | --- | $< 2.37 \times 10^{-4}$ *** |
| *$H3.3^{K36R\text{-}WTR}$* | --- | $< 10^{-6}$ **** | $< 2.37 \times 10^{-4}$ *** | --- |

| **Male** | *$H3.3A^{null}$* | *$H3.3^{K36R}$* | *$H3.3A^{null\text{-}WTR}$* | *$H3.3^{K36R\text{-}WTR}$* |
|---|---|---|---|---|
| *$H3.3A^{null}$* | --- | $< 2.37 \times 10^{-4}$ *** | ns | --- |
| *$H3.3^{K36R}$* | $< 2.37 \times 10^{-4}$ *** | --- | --- | $< 10^{-6}$ **** |
| *$H3.3A^{null\text{-}WTR}$* | ns | --- | --- | ns |
| *$H3.3^{K36R\text{-}WTR}$* | --- | $< 10^{-6}$ **** | ns | --- |

**Figure S3.** Adult longevity assays for *$H3.3A^{null}$* and *$H3.3A^{null\text{-}WTR}$* controls, and for *$H3.3^{K36R}$* and *$H3.3^{K36R\text{-}WTR}$* flies, parsed by sex. Median lifespan was determined (dotted lines) by identifying the day at which 50% of the animals survived. Statistical comparison of survival curves using Gehan-Breslow-Wilcoxon tests are presented in the accompanying table. A Bonferroni correction for multiple comparisons was employed, resulting in the following adjusted significance values: * $p<0.0125$, ** $p<0.0025$, *** $p<2.5 \times 10^{-4}$, ****$p<2.5 \times 10^{-5}$.

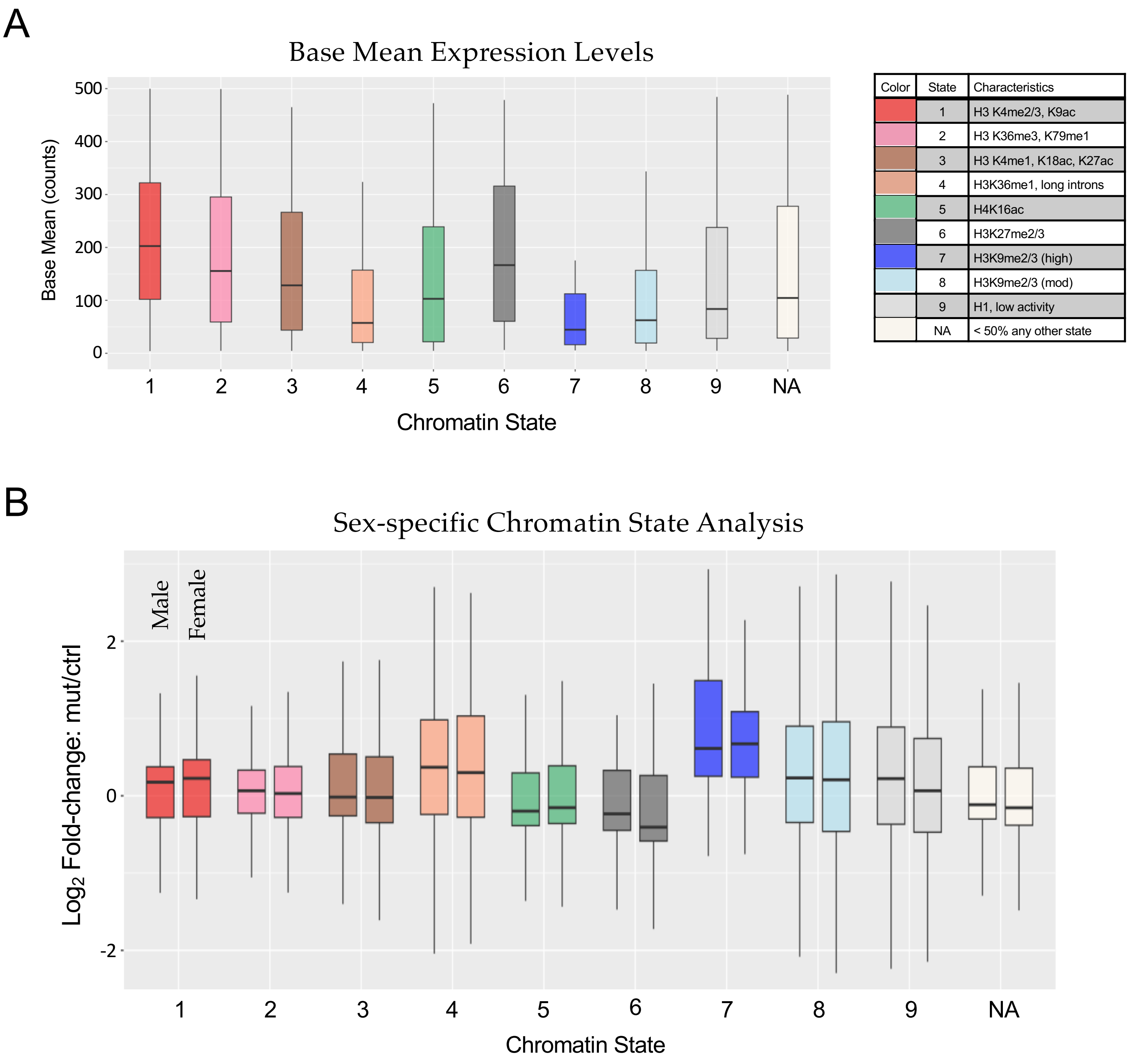


**Figure S4. Additional Chromatin State Analyses**. **A**) The Base Mean gene expression for all genotypes in the non-interaction DESeq2 model was plotted for genes, and binned as in Figure 5A. **B**) For the set of genes described in Figure 5A, and utilizing a DESeq2 model which considers sex as an additional variable (Fig. S2), separate mut/ctrl Log2 Fold-change values were plotted for Old K36R male and female animals.

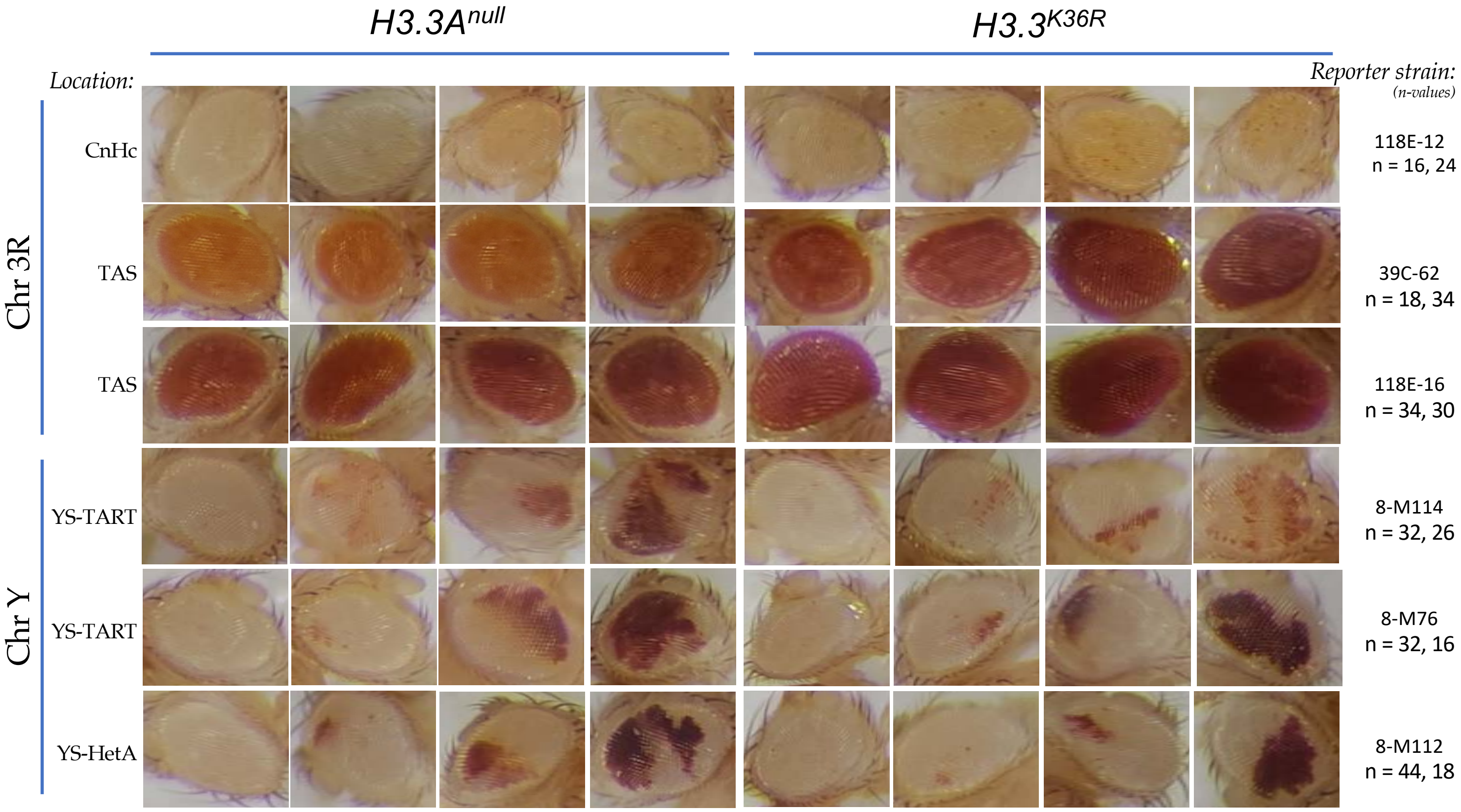


**Figure S5. Representative eyes from Position Effect Reporter strains.** For each row, four representative eyes for a particular reporter transgene in either the *H3.3A*$^{null}$ control or *H3.3*$^{K36R}$ mutant background are shown. To the left, chromosomal location and type of chromatin region where each reporter insertion is located is indicated. Pericentric heterochromatin (CnHc), Telomere Associated Sequences (TAS), YS-TART (Y chromosome TART), and YS-HetA (Y chromosome-HetA) are abbreviated as indicated. The number (n) of eyes evaluated per *H3.3A*$^{null}$ control (ctrl) and *H3.3*$^{K36R}$ mutant (mut) genotype are indicated at right, below the label for each reporter strain and also listed here: 118E-12 (ctrl=16, mut=24), 39C-62 (ctrl=18, mut=34), 118E-16 (ctrl=30, mut=22), 8-M114 (ctrl=32, mut=26), 8-M76 (ctrl=32, mut=16), 8-M112 (ctrl=44, mut=18).
